# SPITROBOT-2 advances time-resolved cryo-trapping crystallography to under 25 ms

**DOI:** 10.1101/2025.05.23.655289

**Authors:** Maria Spiliopoulou, Caitlin E. Hatton, Martin Kollewe, Jan-Philipp Leimkohl, Hendrik Schikora, Friedjof Tellkamp, Pedram Mehrabi, Eike C. Schulz

## Abstract

We previously introduced the SPITROBOT, a protein crystal plunging system that enables reaction quenching via cryo-trapping with a time resolution in the millisecond range. Here we present the next generation, SPITROBOT-2, as an integrated benchtop device, condensed to approximately an A4 footprint. The user experience has been enhanced by the integration of a guiding beam and a sample-switch dial, to optimise sample exchange operations. This is complemented by a light-indicated liquid nitrogen level sensor, ensuring enhanced reliability. Moreover, a fully automated shutter shields the liquid nitrogen from the humidified environment, improving sample integrity. These improvements reduce the net sample preparation time to approximately three minutes per sample. Most importantly, the cryo-trapping delay time has been reduced to 23 milliseconds, making SPITROBOT-2 twice as fast as the previous generation. This further expands the number of target systems that can be addressed by cryo-trapping time-resolved crystallography. We demonstrate successful cryo-trapping via ligand binding and conformational changes using 12 crystal structures of three independent model systems: xylose isomerase, human insulin and bacteriophage T4 lysozyme. Taken together, these improvements increase the convenient access to cryo-trapping, time-resolved X-ray crystallography empowering the MX community with efficient tools to advance research in structural biology.

---

Time-resolved crystallography (TRX) is a powerful technique to study dynamic events and conformational responses of proteins while they carry out their function. In the past decade it has undergone a resurgence, primarily triggered by the advent of X-ray free-electron laser (XFEL) sources and the emergence of serial data collection methods (*1–3*). The successful application of serial crystallography at XFELs was quickly adapted at synchrotron facilities, which can provide equivalent data quality (*4–7*). While serial synchrotron crystallography (SSX) can not tap the ultra-fast time-scales that are accessible to XFELs, they are however, the obvious choice when it comes to studying enzymatic mechanisms beyond the microsecond time domain (*2, 8, 9*). Their comparably wide distribution goes in hand with a lower competition for beamtime and thus enables a more democratic access, connected with the ability to study a larger number of model systems and develop novel methods that will benefit the field as a whole (*6, 9–26*). However, in spite of clear fundamental and practical advantages as well as recent progress in simplifying the method, time-resolved crystallography still requires expert knowledge especially in bridging the divide between crystallography, physics and biochemistry and thus often remains the niche of experts.

To this end cryo-trapping crystallography can display a convenient workaround that permits resolving stable reaction intermediates by quickly quenching biochemical reaction in liquid nitrogen. Traditionally, slow enzymes with turnover times in the seconds domain or those with repetitive mechanisms have been addressed with cryo-trapping for decades (*27, 28*). In addition to its conceptual simplicity, a major advantage of cryo-trapping approaches is rooted in its compatibility with established high-throughput infrastructure and automated data-processing routines. However, a common problem in manual cryo-trapping lies in its limitation to macroscopic crystals, the comparably large time jitter and associated with this, its comparably low reproducibility, especially if time-scales faster than ca. 30 seconds are aimed for. To address these limitations, we have recently developed the *SPITROBOT*, an automatic crystal plunger that enables time-resolved crystallography via cryo-trapping in the millisecond time domain (*29*). Reactions are initiated by the liquid application method for time-resolved applications (LAMA), which permits *in situ* mixing with minimal amount of substrate solution, while allowing for reaction initiation times in the millisecond time-domain (*25*). LAMA nozzles spray picoliter sized droplets of ligand solution onto protein crystals, which are exposed to X-rays or cryo-trapped after a pre-defined delay time. The *SPITROBOT* is compatible with macroscopic crystals, micro-crystals, canonical rotation as well as serial data collection methods. In addition, the *SPITROBOT* is fully compatible with the SPINE standard (e.g. crystals are directly vitrified inside SPINE pucks), which directly connects cryo-trapping to the high-throughput infrastructure available at most synchrotrons. Another major advantage in cryo-trapping approaches lies in the ability to uncouple sample preparation from data collection, that is users can prepare their samples well in advance to a beamtime and fully focus on either task.

For the growing user base interested in cryo-trapping crystallography using the *SPITROBOT*, we have updated the first generation prototype. Now, *SPITROBOT-2* is a fully integrated benchtop device with a minimal footprint conveniently fitting into existing MX-laboratories, even if lab space constraints have to be taken into consideration. Like the first generation *SPITROBOT-2* also maintains humidity and temperature conditions during reaction initiation via implementation of an environmental control system, and thereby also permits addressing long delay times (*29*). Finally, we have substantially advanced the user experience by improving the overall design and workflow of *SPITROBOT-2*. Most importantly, with further hardware improvements we were able to reduce the minimum delay time by a factor of 2, now enabling cryo-trapping within less than 25 ms.

## Results and Discussion

### An integrated benchtop device

In order to turn the *SPITROBOT* prototype into a user-friendly device, we aimed to generate a compact setup with minimal clutter that aids the user in the sample preparation process. For an overview incl. abbreviations please refer to **Fig. 1 a,b**. The footprint of *SPITROBOT-2* was substantially reduced, now comprising dimensions of W284 x H480 x D316 mm, with a weight of approximately 15 kg. Two handles on the sides allow for convenient relocation and enable use as a benchtop device. In contrast to the prototype, *SPITROBOT-2* permits triggering the plunging process from the main device itself, and therefore no longer depends on an external control box. The humidity flow device (HFD) as well as a control computer (for e.g. delay time setup, sample monitoring, automatic image capturing) remain autonomous solutions.

**Figure 1:**
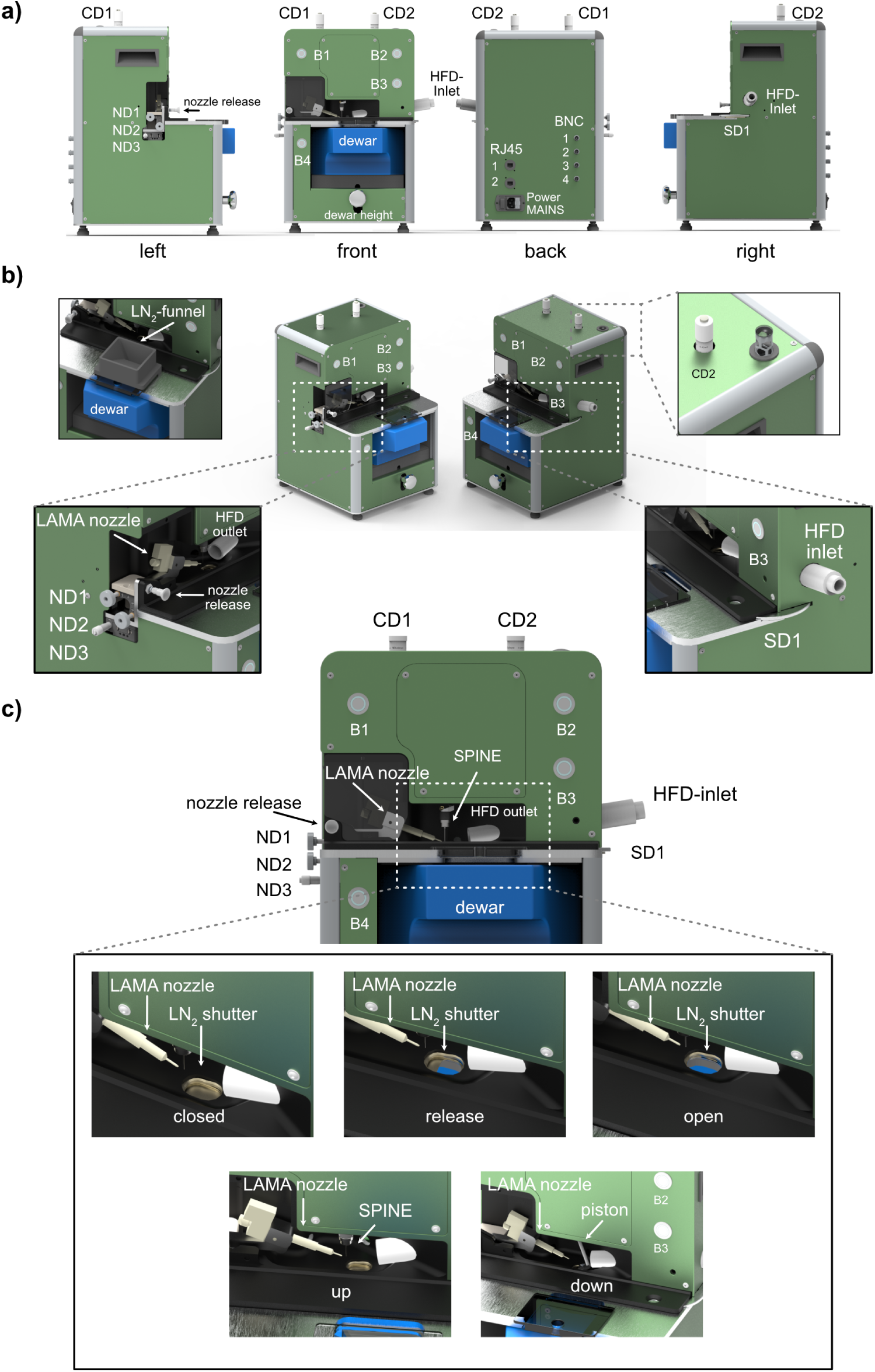
The redesigned *SPITROBOT-2.* a) All side views of *SPITROBOT-2*. b) 2-sided isometric view of *SPITROBOT-2* with closeups of the liquid nitrogen (LN_2_) funnel, the allen key lift, the nozzle dials (ND1-ND3) and the sample dial (SD). c) The front-side of *SPITROBOT-2*, with closeups of the different liquid nitrogen shutter steps and the two main positions of the piston. Custom abbreviations: CD - camera dial; ND - nozzle dial; B - button; SD - sample dial; HFD - humidity flow device.

The front panel comprises the two-hand-control safety switches that were formerly part of the control box (B1, B2). Pressing these buttons triggers reaction initiation and sample plunging according to the settings in the software interface. The two-hand-control solution prevents inadvertent interaction with the sample area during the plunging process and thereby prevents user injury. An additional button (B3) on the right side is used to reset the device in case of an error or as a light switch for sample illumination. To enable high-throughput sample preparation, samples are plunged directly into a SPINE-puck. Like in the prototype, a modified dewar containing the SPINE puck is placed at the bottom and its height is adjusted via an integrated labjack. Sample exchange is executed via a sample exchange dial (SD1), that enables rotating the SPINE puck inside of the dewar to access the next sample position. Next to the dewar, on the lower left of the device another button (B4) can be used to manually release the pin from the electromagnet on the piston. All buttons (B1 - B4) are equipped with a color ring indicating the process or function, e.g. red for error.

The sample viewing system consists of two cameras, aligned at an angle of 90° to each other. On top of *SPITROBOT-2* both cameras can be focused independently via two dials (CD1, CD2). To facilitate non-standard adjustments, such as precise camera centering, the device incorporates a built-in Allen key mechanism, which can be accessed by applying slight pressure to the top of the device. The LAMA nozzle for reaction initiation is mounted from the left side of *SPITROBOT-2* and its precise orientation with respect to the sample position is controlled via three nozzle dials (ND1, ND2, ND3). The nozzle can be locked in a retracted position for sample mounting and quickly released into spraying position by the nozzle release latch. Furthermore, the nozzle can be replaced by focusing optics, thus enabling optical excitation of the sample by a fibre-coupled light source or laser (*REF: Mehrabi, Standfuss et al. - in press*). On the right side of *SPITROBOT-2* the HFD-nozzle is fed through to the sample position. On the back, there are two network ports, four BNC connections (e.g. for the LAMA trigger signal) as well as the main power supply and switch.

### A liquid nitrogen level control system ensures sample quality

Sample quality is key to any TRX experiments, but for cryo-trapping TRX this also includes the quality of the cryogen. The quality of the vitrified samples critically depends on the steepness of the temperature gradient and the purity of the liquid nitrogen, ie its level of water-ice contamination (*30–34*). To support this, integrated temperature sensors will warn the user or even throw an error if the level of liquid nitrogen drops too low, which would compromise the vitrification process. Liquid nitrogen refilling is guided by a safety funnel, that enables refilling while the dewar remains in the sample preparation position. To insulate the liquid nitrogen from the atmospheric air and the stream of highly humid warm air from the HFD, *SPITROBOT-2* is equipped with a liquid nitrogen shutter window, that only opens during the plunging period (**Fig. 1 c**) but shuts access to the cryogen at all other times. This limits liquid nitrogen evaporation and thus increases the cryogen life time in the dewar, but more importantly it reduces ice contamination and thereby improves sample quality.

There is a clear influence of plunge freezing speeds and sample volumes on cooling rates and vitrification outcomes, providing, thus, insights for further optimization. Previous studies on cryofixation emphasize that eliminating the cold gas layer during plunge freezing enhances cooling rates improving sample preservation (*35*). Most of the cooling occurs in the cold gas layer above the cryogen during plunge cooling of protein crystals in liquid nitrogen (*33*) and for plunge speeds of 1 *ms*^−1^, a 2 cm gas layer is sufficient to dominate cooling (*33,36*). *SPITROBOT-2* is capable of efficient plunge freezing at 2 *ms*^−1^, resulting in further mitigation of the aforementioned phenomenon.

### *SPITROBOT-2* enables delay times of under 25 ms

Early vitrification experiments have provided valuable information on the cooling rates required to achieve successful vitrification. For pure water, these need to be as high as 10^6^*Ks*^−1^, but can be reduced by orders of magnitude by the presence of solutes (*30, 37*). Initial studies using liquid propane near its melting point as a cryogen achieved a peak cooling rate of about 370,000 *Ks*^−1^ with 25-75 µm thermocouples immersed at a rate of 2 *ms*^−1^ (*38*). Further research has highlighted significant differences in the cooling rates of different cryogenic media. In particular, the cooling rates in boiling liquid nitrogen have been reported to be about 50 times lower than those in, for example, 90 K ethane (*39*). Although liquid nitrogen requires the use of cryo-protectants, the reduced safety concerns compared to flammable cryogens have made it the primary cryogen for X-ray crystallography (*31, 40*) and thus the primary choice for the *SPITROBOT-2*.

The electro-pneumatic piston of the prototype has been replaced by a motorised solution, which in *SPITROBOT-2* drives the sample into a vial submerged in liquid nitrogen. In total, it is equipped with two linear motors (Faulhaber LM series), the first of which is used for the plunging process, while the second controls a shutter between the dispensing chamber and the Dewar. The plunger drives (**Fig. 1 c**) the samples into the cryogen at an average speed of 2 *ms*^−1^ (50 mm in 25 ms). To experimentally validate the minimal delay time that can be achieved with *SPITROBOT-2*, we measured the temperature evolution by plunging a 13 µm thermocouple into liquid nitrogen, with and without applying the humidity stream, as described previously (*29*). As these data clearly show, *SPITROBOT-2* allows complete vitrification of the samples within 23 ms, which is about twice as fast as the prototype (**Fig. 2 a**). For crystal sizes that are typically still useful for diffraction experiments at synchrotron sources (about 10-20 µm), this delay time now approaches the theoretical lower diffusion time limit for fast diffusing ligands such as glucose (15 ms) (*9, 25, 41*).

**Figure 2:**
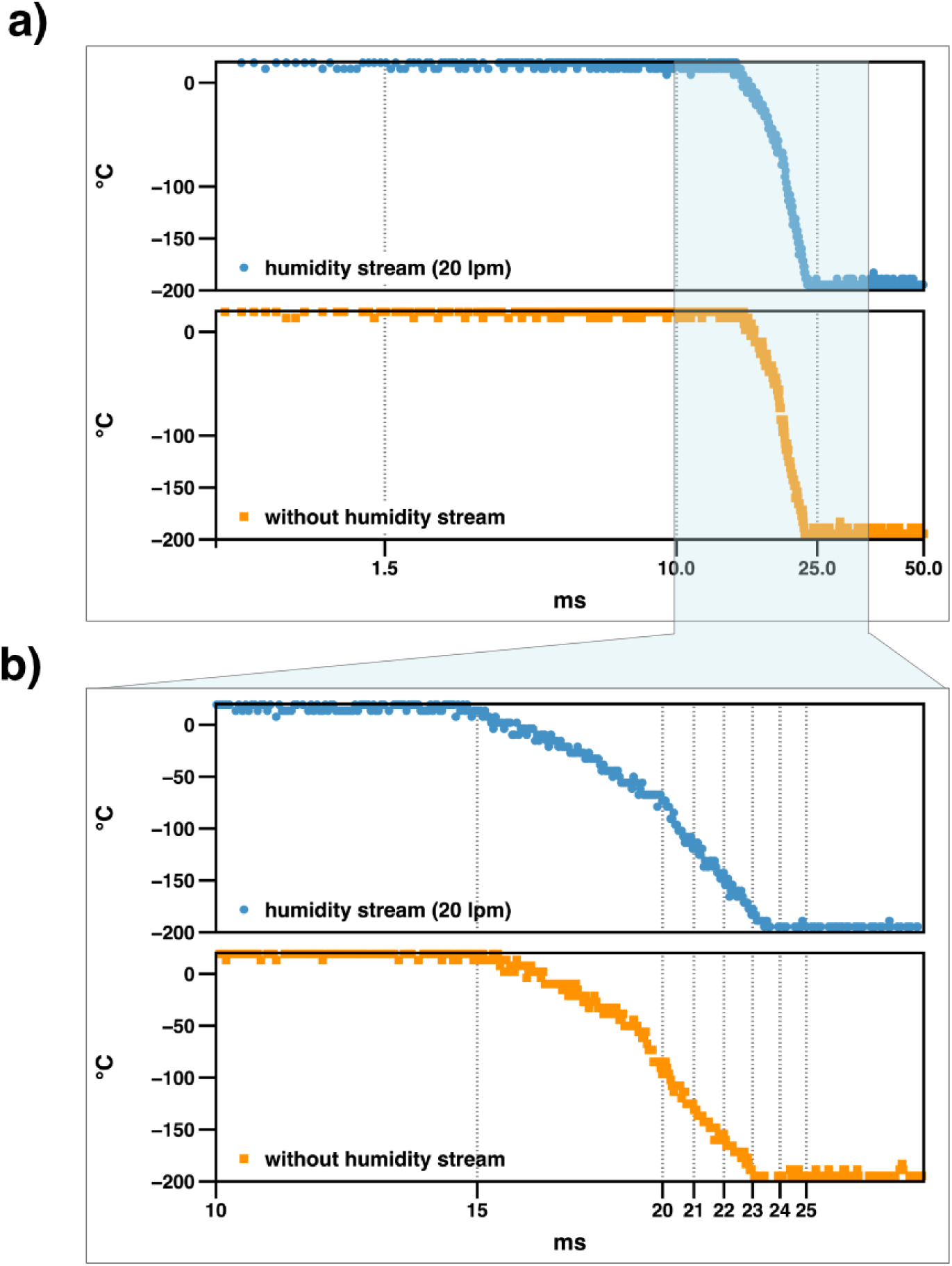
*SPITROBOT-2* vitrification time in liquid nitrogen. a) A global view of the temperature evolution between 0.5 and 50 ms. Irrespective of the humidity flow device the temperature drops to below -196°C, within less than 25 ms. b) closeup of the vitrification time between 10 and 30 ms demonstrates that the vitrification process in liquid nitrogen takes around 7.5 ms and thus matches the vitrification time previously reported for the first generation.

Notably, the vitrification time itself is unaffected by the hardware modification and remains within a time scale of 7.5 ms (**Fig. 2 b**). This means that by immersing samples in liquid nitrogen, protein microcrystals can be vitrified at a cooling rate of approximately 30,000 *Ks*^−1^. We anticipate that by selecting alternative cryogenic media, higher cooling rates may be possible with *SPITROBOT-2* in the future, potentially bypassing the need for cryo-protection.

### Sample preparation workflow

The original workflow of the *SPITROBOT* has been redesigned to provide a more user-friendly, streamlined experience that speeds up the entire sample preparation process. To start a sample preparation, the pre-cooled dewar containing the SPINE puck is now placed on an integrated lifting mechanism. Once the dewar has been inserted, precise positioning within the *SPITROBOT-2* is aided by a guide beam which helps the user to align the puck in relation to the rotation axis of the instrument.

Once in position, the dewar is lifted by the labjack until it seals against the insulated and actively heated plate separating the dewar from the upper part of the machine. This has two advantages: firstly, it helps to protect the operator during sample vitrification; secondly, it also greatly reduces ice formation within the liquid nitrogen as the dewar is now isolated from both the atmosphere and the moisture stream. Sample access to the liquid nitrogen is only possible via the integrated shutter (*see above*).

Next, the semi-manual sample dial on the right-hand side of the *SPITROBOT-2* is used to position the start vial in the immersion position. The alignment of the dispenser nozzle is supported by an integrated dual camera system (IDS imaging). Now the desired delay time can be set in the control software and the system is ready to start the plunging process by pressing the two-hand-control safety switches. Whilst retaining the two-handed safety switch solution, these have been integrated into the main unit so that *SPITROBOT-2* is no longer dependent on an external control box. Moreover, colour-coded signals integrated into the control buttons provide the user with immediate feedback about the plunging process, e.g. green and blue to let the operator know when to press or release buttons, or a white flash at the moment of sample excitation.

### Ligand binding events and conformational changes

To demonstrate the versatile applicability of *SPITROBOT-2* we demonstrate reaction initiation and fast delay times in three independent model systems, obtained by single crystal rotation data collection: xylose isomerase (XI), human insulin (HI) and bacteriophage T4 lysozyme (T4L).

### Xylose isomerase ligand binding trapped in 25 ms

As a direct comparison to the first generation of *SPITROBOT*, we use xylose isomerase (XI) as a model system. XI is of great commercial interest as it catalyses the reversible interconversion of *α*-D-glucose to *β*-D-fructose (*42, 43*). We have previously shown that glucose can bind to XI within 15 ms, which approximates theoretical diffusion times for crystals of this size (*9, 25, 41*). To initiate substrate binding, we sprayed a 2 M glucose solution onto 20 µm XI crystals loaded on a micromesh before plunging them in liquid nitrogen with delay times of 25 ms and 50 ms, respectively. The difference density map in the active site shows that with *SPITROBOT-2* we can capture the diffusion of glucose after the minimum delay time of 25 ms (**Fig. 3**). This means that our current data confirm previous observations on glucose diffusion kinetics in protein crystals (*25*).

**Figure 3:**
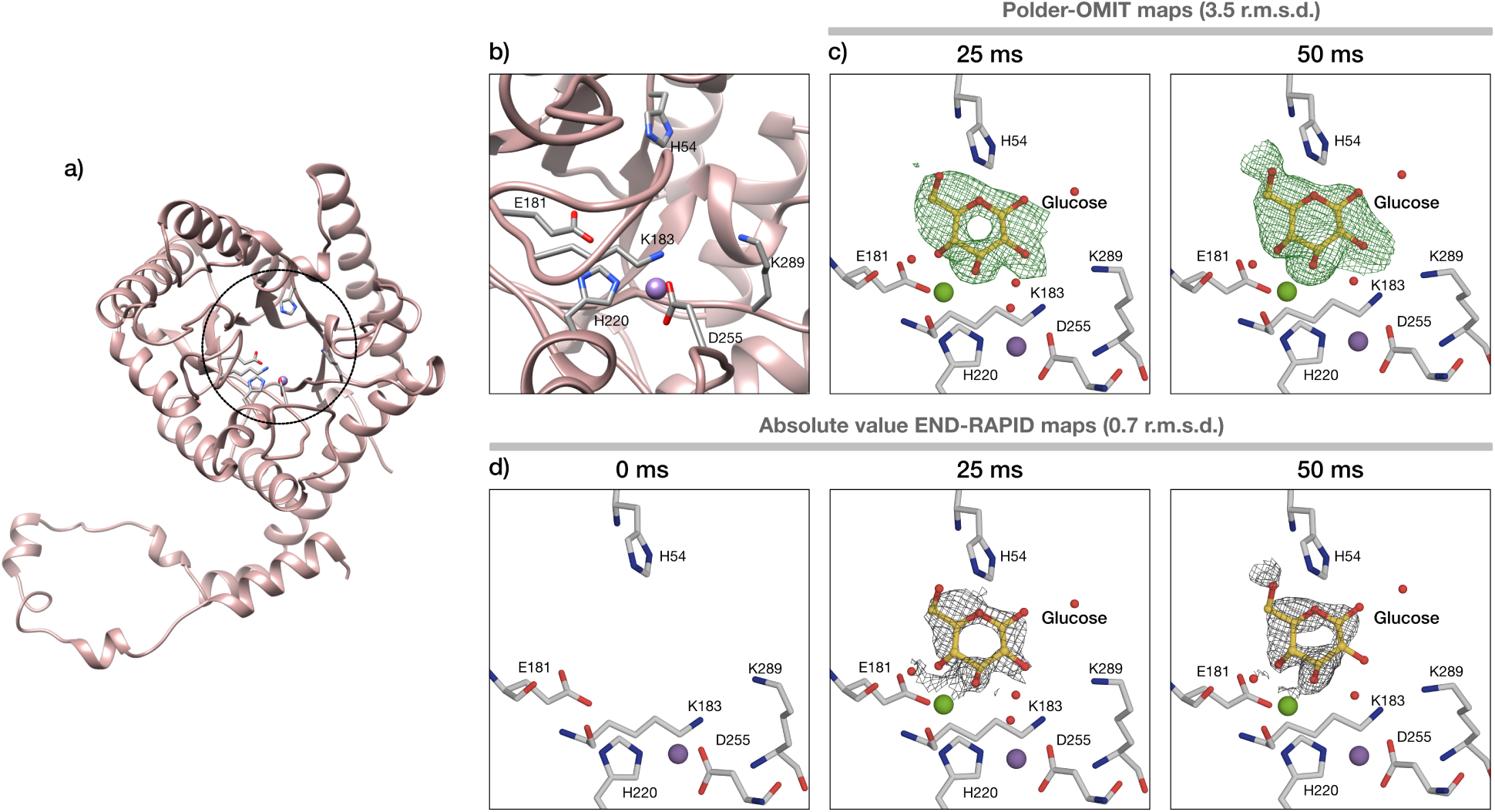
Glucose binding in the active site of XI. a) Cartoon representation of XI structure; the active site is indicated by a circle. b) Closeup of the active site and surrounding residues. c, d) Polder-OMIT maps and absolute value END-RAPID maps for glucose binding after 25 and 50 ms of soaking.

### A pH jump triggers conformational changes in human insulin

Human insulin (HI), is a 5.8 kDa peptide hormone produced by *β*-pancreatic cells, which facilitates the absorption of carbohydrates from the blood into tissues. It was one of the first proteins to be isolated and crystallographically studied (*44, 45*). In its active form, HI consists of 51 amino acids divided into two polypeptide chains: A and B (21 and 30 amino acids respectively). The secondary structure of insulin shows two almost antiparallel *α*-helices in chain A and a *α*-helix followed by a turn and a *β*-strand in chain B (*46*). After a century of crystallographic research, insulin has been identified as one of the most polymorphic proteins, displaying a range of molecular conformations, crystal forms and binding capacities for metals and ligands (*47*). Several cubic insulin crystal structures have been analysed under different conditions, such as crystals in 0.1 M sodium salt solutions with pH values ranging from 7.0 to 10.0 (*48*) and in 1 M Na_2_SO_4_ at pH 5.0 to 11.0 (*49*). It has previously been reported that cubic insulin crystals undergo a structural transition when the pH is lowered from 9.0 to 5.0 (*49*). The conformational change of glutamate at position 13 of chain B (GluB13) is accompanied by the binding of sulphate ions close to phenylalanine at position 1 of chain B (PheB1). Here we aimed to induce this effect by a pH jump with *SPITROBOT-2*.

To this end, we sprayed a low pH sulphate buffer (1 M CH_3_COONa, pH 4.5, 1 M Na_2_SO_4_, 15% (v/v) ethylene glycol) onto the insulin crystals and followed the transition of GluB13 and SO_4_ binding as a function of time. As can be clearly seen from the difference density maps, both the structural transition and the SO_4_ binding can be observed at delay times as short as 25 ms (**Fig. 4**; for refined occupancies for GluB13 and SO_4_ see **Table S2**).

**Figure 4:**
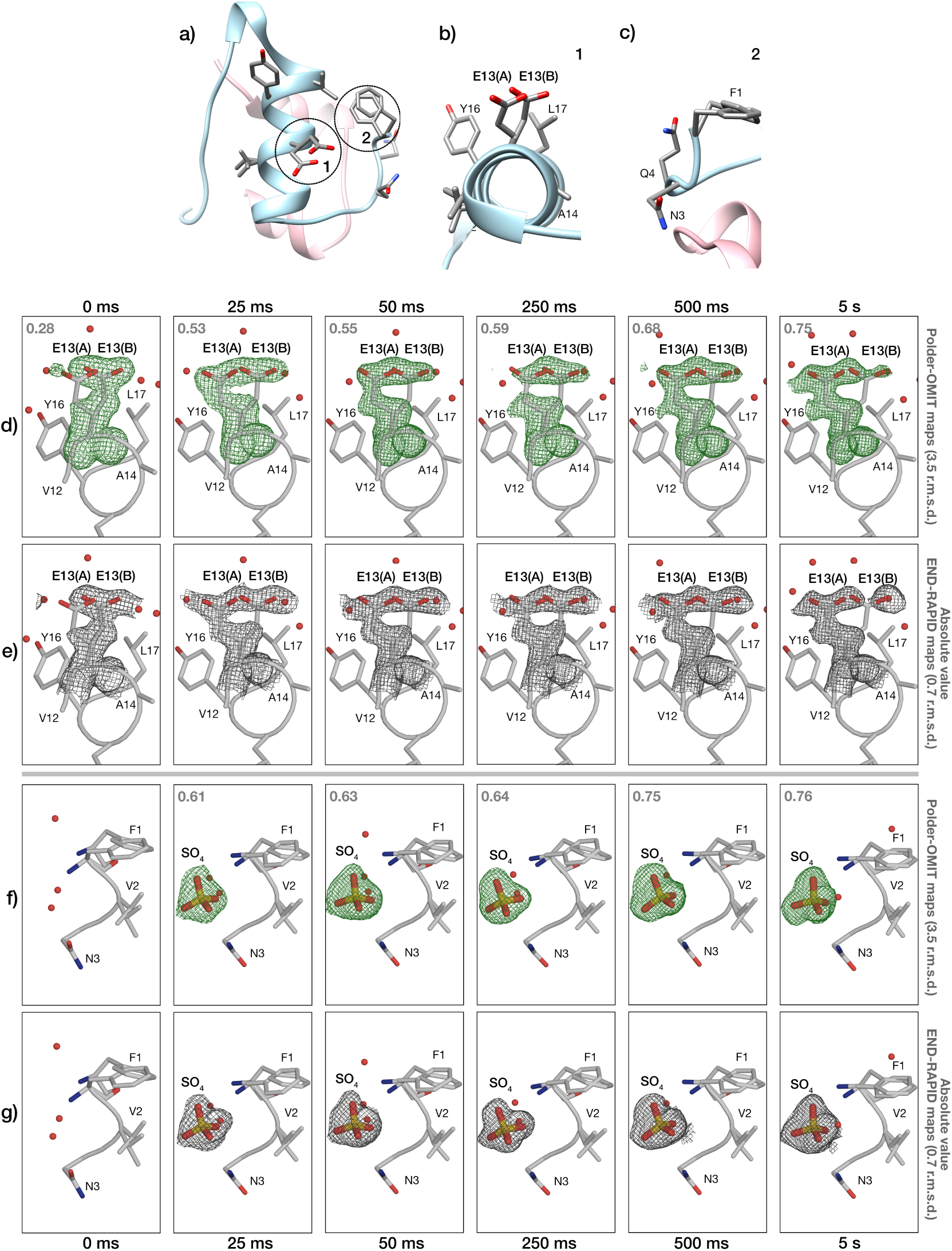
pH-jump effect on HI. a) Cartoon representation of HI structure; the two chains are illustrated in pink (chain A) and blue (chain B), while the residues in the GluB13 region (1) and the SO_4_ binding region (2) are represented in sticks. b) Closeup of GluB13 and surrounding residues for HI at pH 9.0. c) Closeup of the SO_4_ binding region at pH 9.0 (apo state). d, e) Polder-OMIT maps and absolute value END-RAPID maps for the effect of pH 4.5 on the GluB13 after 25 ms to 5 s; the occupancies of conformation A are indicated in grey. f, g) Polder-OMIT maps and absolute value END-RAPID maps for the effect of pH 4.5 on SO_4_ binding after 25 ms to 5 s soaking; the occupancies for SO_4_ are indicated in grey.

### Backbone changes in bacteriophage T4-lysozyme (T4L)

The biological function of T4L is to hydrolyse the *β*-1,4 linkage between N-acetylmuramic acid and N-acetylglucosamine in bacterial peptidoglycan to aid cell lysis (*50,51*). The L99A mutation of T4L forms an internal non-polar cavity of about 150 Å^3^, which allows the binding of various ligands after large conformational changes that open T4L to allow ligand binding (*52, 53*). As a result, this particular mutant serves as a model system for studying ligand binding at allosteric sites. Previous studies focusing on benzene binding pathways have proposed that structural rearrangements of the cavity surrounding the helices allow the ligand to enter the binding site (*52–54*). To explore whether we could induce and capture larger conformational changes, we used indole as a ligand and observed structural changes after 1 s and 10 s, respectively. Our *SPITROBOT-2* cryo-trapping data clearly show that structural snapshots of the binding event can be obtained at delay times (1 s) inaccessible by manual methods (**Fig. 5**). In addition to indole binding, we clearly observe a conformational change in the backbone of the *F*-helix of T4L in the indole-bound states, which is shifted by about 1.8 Å compared to the apo state (**Fig. 5e**).

**Figure 5:**
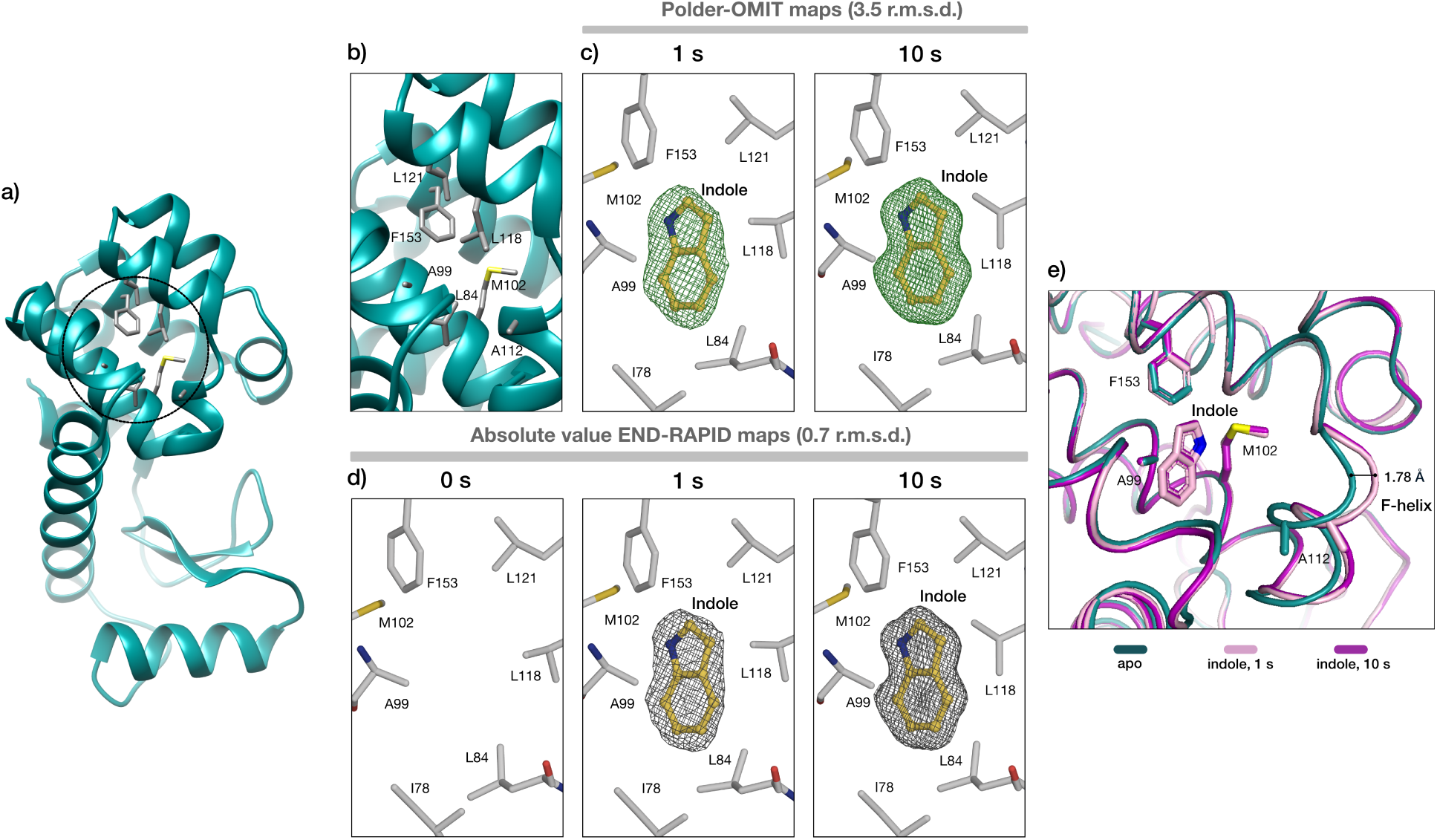
Indole binding in the non-polar cavity of T4L-L99A. a) Cartoon representation of T4L-L99A structure; the cavity is indicated by a circle. b) Closeup of the cavity and the surrounding residues. c, d) Polder-OMIT maps and absolute value END-RAPID maps for indole binding after 1 s and 10 s of soaking time. e) T4L-L99A cavity closeup. Ribbon representation of protein cavity in the absence (teal) and presence of indole after 1 s (pink) and 10 s (magenta) soaking time. The conformational change of the *F*-helix can be observed for the bound states of T4L, leading to a further opening of the cavity (1.78 Å) in the presence of the ligand.

## Conclusions

*SPITROBOT-2* is an integrated benchtop solution for cryo-trapping protein crystals with a delay time of less than 25 ms. It is suitable for monitoring ligand binding events as well as conformational changes of main and side chains. As the majority of enzymes have a turnover number of around 10 s^−1^ (*55*), a large number of systems should be amenable to cryo-trapping TRX, e.g. to track catalytic intermediates that are inaccessible by conventional methods. Its small footprint, versatility and conceptual simplicity should make *SPITROBOT-2* an attractive asset for MX labs wishing to standardize their vitrification process or explore cryo-trapping TRX. Its strength is further emphasised by its full compatibility with the high throughput workflows found on most MX beamlines.

## Materials and methods

### Temperature measurements

To assess the temperature evolution during the plunging process we attached a 13 µm type K thermocouple (KFT-13-200-200, As One Corporation) to a canonical SPINE pin. This modified test sample was plunged into liquid nitrogen with and without application of the humidity flow device. Temperature corresponding voltage measurements were recorded in 50 µs intervals.

### Protein purification and crystallization

#### Xylose isomerase

Xylose isomerase in pET24a vector (Genscript), was expressed in BL21 DE3 *E. coli* cells, purified by affinity (Ni) and size exclusion chromatography with a final buffer of 0.05 M Tris pH 8.5, 0.15 M NaCl, with crystallisation as reported in previous studies (*25*). Briefly: 80 mg/mL XI was combined with an equal volume of crystallization buffer (35% (w/v) PEG 3350, 0.2 M LiSO_4_ and 0.01 M HEPES/NaOH, pH 7.5) via SpeedVac crystallization as previously described (*56*).

#### Human insulin

Human insulin purchased by Roche (LOT: 70272900) was diluted in 0.05 M Na_2_HPO_4_, 0.001 M Na_2_EDTA, pH 11.0 (adjusted) to a final concentration of 30 mg/mL. Batch crystallization was performed in a total volume of 160 µL my mixing equal volumes of HI and crystallization buffer consisting of 1.0 M NaH_2_PO_4_/K_2_HPO_4_ pH 9.0 (*57*).

#### Bacteriophage T4 lysozyme

T4-L99A mutant lysozyme gene was cloned in pET29b(+) vector (Genscript) and expressed in BL21 DE3 *E. coli* cells. The protein was purified by ion exchange and size exclusion chromatography with a final buffer of 0.05 M NaH_2_PO_4_/Na_2_HPO_4_ pH 5.5, 0.1 M NaCl, 0.002 M EDTA (*58*). Batch crystallization was performed in a total volume of 20 µL by mixing equal volumes of protein at 22 mg/mL and crystallization buffer consisting of 4.0 M NaH_2_PO_4_/K_2_HPO_4_ pH 7.0, 0.1 M *1,6*-hexanediol and 0.15 M NaCl (PDB ID: 3K2R).

### *SPITROBOT-2* experiment parameters

Crystals of the three individual protein systems were used for the *SPITROBOT-2* experiments. The conditions and solutions used in each case are mentioned in **Table 1**.

**Table 1:** *SPITROBOT-2* samples and environmental details.

| Sample | XI | HI | T4L-L99A |
| --- | --- | --- | --- |
| Crystal size ($\mu\text{m}$) | 20 | 20 | 20 |
| Temperature ($^{\circ}\text{C}$) | 25 | 25 | 25 |
| Relative humidity (%) | 95 | 95 | 95 |
| Ligand | Glucose | $\text{Na}_2\text{SO}_4$ | Indole |
| LAMA solution | 0.01 M HEPES pH 7.5, 0.1 M $\text{LiSO}_4$, 0.1 M $\text{MgCl}_2$, 2 M glucose | 1 M $\text{CH}_3\text{COONa}$ pH 4.5, 1 M $\text{Na}_2\text{SO}_4$, 15% (v/v) ethylene glycol | 2.0 M $\text{NaH}_2\text{PO}_4/\text{K}_2\text{HPO}_4$, pH 7.0, 0.15 M NaCl, 0.1 M 1,6-hexanediol, 6% (v/v) DMSO, 0.05 M indole, 20% (v/v) glycerol |

### X-ray data collection

Single crystal rotation datasets were collected at beamline P14 of PETRA-III (DESY, Hamburg) at the EMBL unit Hamburg as well as at beamline ID30A-3 of ESRF (Grenoble, France). Data were collected on an Eiger2 CdTe 16 M detector at 12.7 keV and an Eiger1 X 4M detector at 12.81 keV respectively for the aforementioned beamlines. Data collection parameters for each individual protein case are summarized in **Tables S1, S2 and S3** for XI, HI and T4L-L99A correspondingly.

### Data processing

Diffraction data were automatically processed with autoPROC using StarAniso (*59–62*). Molecular replacement was conducted in Phaser using PDB-ID 8AWS, 1B18 and 4W51, respectively as starting models for XI, HI and T4L-L99A (*63*). Structures were refined using iterative cycles of REFMAC or phenix.refine and coot (*64–66*). Molecular images were generated in PyMol (*67*).

## Funding

The authors gratefully acknowledge the support provided by the Max Planck Society. PM acknowledges support from the Deutsche Forschungsgemeinschaft (DFG) via grant No. 451079909 and from a Joachim Herz Stiftung add-on fellowship. ES acknowledges support by the DFG via grant No. 458246365, and by the Federal Ministry of Education and Research, Germany, under grant number 01KI2114. Funded by the European Union. Views and opinions expressed are however those of the author(s) only and do not necessarily reflect those of the European Union or the European Research Council Executive Agency (ERCEA). Neither the European Union nor the granting authority can be held responsible for them.

## Author contributions

M.S. performed protein expression, purification, prepared the protein crystals, performed data collection, analyzed the data and wrote the manuscript; C.E.H. performed protein expression, purification, crystallization, data collection and analysis; M.K, J.-P.L., H.S. and F.T. designed and validated the spitrobot-2 and the humidity flow device (HFD) and designed the LabView interfaces; F.T., P.M. and E.C.S. designed the experiment and conceptualized the spitrobot-2; E.C.S. supervised and wrote the manuscript; all authors discussed and corrected the manuscript.

## Competing interests

There are no competing interests to declare.

## Data and materials availability

The data that support this study are available from the corresponding authors upon request. All crystallographic data have been deposited in the Protein Data Bank (PDB). Further details are available in Supplementary Tables S1-S3.

## Supplementary Materials for

### Supplementary Text

#### SPINE- to UniPuck sample transfer

A common issue raised by external users of the prototype was the missing compatibility with UniPuck containers (https://smb.slac.stanford.edu/robosync/Universal_Puck/).

Although there is a commercial solution, the CombiPuck™ from MiTeGen, which enables sending samples stored in SPINE standard vials to beamlines that require UniPucks. For transferring samples between SPINE and UniPucks we provide details for in-house manufacturing of a similar transfer solution. We believe that this will alleviate previous concerns and simplify sample transfer between the different puck standards.

#### Data collection and refinement statistics

**Table S1:** Data collection and refinement statistics of XI data. *Values in the highest resolution shell are shown in parentheses*.

| Sample | XI (apo) | XI (glucose, 25 ms) | XI (glucose, 50 ms) |
| --- | --- | --- | --- |
| PDB ID | 9R45 | 9R46 | 9R47 |
| **Data collection** |  |  |  |
| Source | P14, EMBL | P14, EMBL | P14, EMBL |
| Beam size (μm) | 7 x 3 | 7 x 3 | 7 x 3 |
| Flux (ph/s) | 9.254 x 10<sup>11</sup> | 2.037 x 10<sup>11</sup> | 2.037 x 10<sup>11</sup> |
| Energy (keV) | 12.7 | 12.7 | 12.7 |
| Exposure time (ms) | 7.5 | 7.5 | 7.5 |
| Number of images | 3,600 | 10,800 | 3,600 |
| Space group | *I*222 | *I*222 | *I*222 |
| Unit cell parameters |  |  |  |
| *a*, *b*, *c* (Å) | 85.498, 93.723, 98.033 | 85.680, 93.760, 98.070 | 85.643, 93.922, 98.322 |
| $\alpha$, $\beta$, $\gamma$ (°) | 90 | 90 | 90 |
| Resolution range (Å) | 67.745-1.788 (2.018-1.788) | 63.250-2.100 (2.175-2.10) | 67.915-1.802 (1.948-1.802) |
| Total reflections | 141,351 (6,541) | 884,954 (39,157) | 401,901 (17,336) |
| Unique reflections | 20,717 (977) | 23,420 (1692) | 30,012 (1,501) |
| Mean *I*/$\sigma$(*I*) | 5.9 (1.6) | 24.6 (6.1) | 8.4 (1.5) |
| Completeness (ellipsoidal) | 88.3 (50.9) | 100 (100) | 93.8 (54.7) |
| Multiplicity | 6.8 (6.7) | 37.8 (24.0) | 13.4 (11.5) |
| CC<sub>1/2</sub> | 0.983 (0.547) | 0.999 (0.958) | 0.996 (0.464) |
| **Refinement** |  |  |  |
| Resolution range | 67.74-1.79 | 64.52-2.10 | 49.16-1.80 |
| Number of reflections | 20,713 | 23,389 | 30,001 |
| *R*<sub>work</sub> | 19.57 | 18.69 | 16.92 |
| *R*<sub>free</sub> | 24.44 | 24.26 | 20.65 |
| Occupancy - Glucose | - | 0.53 | 0.56 |
| Wilson B-factor (Å<sup>2</sup>) | 18.56 | 27.19 | 21.80 |
| Mean B-factor (Å<sup>2</sup>) |  |  |  |
| Overall | 20.70 | 29.49 | 23.75 |
| Protein | 20.31 | 29.12 | 22.97 |
| Water | 26.06 | 33.63 | 30.68 |
| Other | 41.85 | 33.17 | 29.41 |
| **Model quality** |  |  |  |
| *RMS deviations* |  |  |  |
| Bond length (Å) | 0.007 | 0.007 | 0.007 |
| Bond angles (°) | 0.874 | 0.850 | 0.834 |

**Table S2:** Data collection and refinement statistics of HI data. *Values in the highest resolution shell are shown in parentheses*.

| Sample | HI (pH 9.0, apo) | HI (pH 4.5, 25 ms) | HI (pH 4.5, 50 ms) | HI (pH 4.5, 250 ms) | HI (pH 4.5, 500 ms) | HI (pH 4.5, 5 s) |
| --- | --- | --- | --- | --- | --- | --- |
| PDB ID | 9R48 | 9R49 | 9R4A | 9R4B | 9R4C | 9R4E |
| **Data collection** |  |  |  |  |  |  |
| Source | ID30A-3, ESRF | ID30A-3, ESRF | ID30A-3, ESRF | ID30A-3, ESRF | ID30A-3, ESRF | ID30A-3, ESRF |
| Beam size (μm) | 15 x 15 | 15 x 15 | 15 x 15 | 15 x 15 | 15 x 15 | 15 x 15 |
| Flux (ph/s) | 5.22 x 10<sup>11</sup> | 5.18 x 10<sup>11</sup> | 5.23 x 10<sup>11</sup> | 5.16 x 10<sup>11</sup> | 5.23 x 10<sup>11</sup> | 5.20 x 10<sup>11</sup> |
| Energy (keV) | 12.8 | 12.8 | 12.8 | 12.8 | 12.8 | 12.8 |
| Exposure time (ms) | 10 | 10 | 10 | 10 | 10 | 10 |
| Number of images | 900 | 900 | 900 | 900 | 900 | 900 |
| Space Group | *I*2<sub>1</sub>3 | *I*2<sub>1</sub>3 | *I*2<sub>1</sub>3 | *I*2<sub>1</sub>3 | *I*2<sub>1</sub>3 | *I*2<sub>1</sub>3 |
| Unit cell parameters |  |  |  |  |  |  |
| *a* (Å) | 78.802 | 78.396 | 78.345 | 78.398 | 78.438 | 78.131 |
| *α* (°) | 90 | 90 | 90 | 90 | 90 | 90 |
| Resolution range (Å) | 39.401-1.650<br>(1.736-1.650) | 39.198-1.641<br>(1.713-1.641) | 39.173-1.439<br>(1.549-1.439) | 39.199-1.493<br>(1.572-1.493) | 39.219-1.407<br>(1.486-1.407) | 39.065-1.293<br>(1.387-1.293) |
| Total reflections | 71,234 (3,818) | 59,591 (3,198) | 72,904 (2,577) | 115,848 (6,117) | 111,825 (6,000) | 90,466 (5,166) |
| Unique reflections | 8,996 (450) | 9,036 (452) | 12,209 (610) | 11,791 (590) | 13,933 (697) | 16,218 (811) |
| Mean *I*/*σ*(*I*) | 9.9 (1.3) | 6.4 (1.3) | 8.2 (1.3) | 8.2 (1.6) | 8.9 (1.4) | 6.5 (1.4) |
| Completeness (ellipsoidal) | 94.1 (44.6) | 93.3 (49.4) | 94.4 (40.5) | 94.2 (48.3) | 94.1 (47.1) | 91.4 (41.9) |
| Multiplicity | 7.9 (8.5) | 6.6 (7.1) | 6.0 (4.2) | 9.8 (10.4) | 8.0 (8.6) | 5.6 (6.4) |
| CC<sub>1/2</sub> | 0.996 (0.541) | 0.982 (0.313) | 0.994 (0.387) | 0.994 (0.335) | 0.997 (0.424) | 0.992 (0.319) |
| **Refinement** |  |  |  |  |  |  |
| Resolution range | 39.43-1.65 | 39.23-1.64 | 39.20-1.45 | 39.23-1.50 | 39.25-1.41 | 39.10-1.30 |
| Number of reflections | 8,539 | 8,594 | 11,584 | 11,187 | 13,225 | 15,412 |
| *R*<sub>work</sub> | 18.604 | 17.916 | 17.041 | 18.047 | 17.308 | 17.586 |
| *R*<sub>free</sub> | 19.625 | 19.760 | 19.565 | 20.824 | 20.062 | 19.370 |
| Occupancy - GluB13 (A/B) | 0.28/0.72 | 0.53/0.47 | 0.55/0.45 | 0.59/0.41 | 0.68/0.32 | 0.75/0.25 |
| Occupancy - SO<sub>4</sub> | - | 0.61 | 0.63 | 0.64 | 0.75 | 0.76 |
| Wilson B-factor (Å<sup>2</sup>) | 16.41 | 10.66 | 12.29 | 12.13 | 10.75 | 10.16 |
| Mean B-factor (Å<sup>2</sup>) |  |  |  |  |  |  |
| Overall | 23.45 | 16.77 | 18.84 | 18.12 | 16.57 | 15.02 |
| Protein | 23.13 | 16.16 | 21.37 | 17.61 | 15.78 | 14.58 |
| Water | 34.80 | 29.50 | 31.20 | 31.50 | 30.30 | 26.20 |
| Other | — | 32.50 | 31.00 | 27.30 | 28.20 | 19.00 |
| **Model quality** |  |  |  |  |  |  |
| *RMS deviations* |  |  |  |  |  |  |
| Bond length (Å) | 0.0115 | 0.0124 | 0.0161 | 0.0136 | 0.0132 | 0.0130 |
| Bond angles (°) | 2.05 | 1.94 | 2.22 | 1.97 | 1.82 | 1.88 |

**Table S3:** Data collection and refinement statistics of T4L-L99A data. *Values in the highest resolution shell are shown in parentheses*.

| Sample | T4L-L99A (apo) | T4L-L99A (indole, 1 s) | T4L-L99A (indole, 10 s) |
| --- | --- | --- | --- |
| PDB ID | 9R4F | 9R4G | 9R4H |
| **Data collection** |  |  |  |
| Source | P14, EMBL | P14, EMBL | P14, EMBL |
| Beam size ($\mu\text{m}$) | 50 x 50 | 50 x 50 | 30 x 30 |
| Flux (ph/s) | $1.171 \times 10^{12}$ | $4.208 \times 10^{11}$ | $1.441 \times 10^{12}$ |
| Energy (keV) | 12.7 | 12.7 | 12.7 |
| Exposure time (ms) | 7.5 | 7.5 | 7.5 |
| Number of images | 3,600 | 3,600 | 3,600 |
| Space group | $P3_221$ | $P3_221$ | $P3_221$ |
| Unit cell parameters |  |  |  |
| $a, b, c$ ($\text{\AA}$) | 60.097, 60.097, 96.251 | 60.173, 60.173, 97.217 | 60.241, 60.241, 97.706 |
| $\alpha, \beta, \gamma$ ($^\circ$) | 90, 90, 120 | 90, 90, 120 | 90, 90, 120 |
| Resolution range ($\text{\AA}$) | 52.046-1.497 (1.567-1.497) | 52.111-1.896 (2.008-1.896) | 52.170-1.493 (1.560-1.493) |
| Total reflections | 550,310 (28,073) | 214,047 (10,832) | 591,509 (29,839) |
| Unique reflections | 29,721 (1,383) | 11,696 (586) | 31,276 (1,548) |
| Mean $I/\sigma(I)$ | 12.4 (1.5) | 8.4 (1.3) | 15.9 (1.4) |
| Completeness (ellipsoidal) | 94.5 (51.9) | 87.4 (51.4) | 95.1 (50.5) |
| Multiplicity | 18.5 (20.3) | 18.3 (18.5) | 18.9 (19.3) |
| CC<sub>1/2</sub> | 0.998 (0.525) | 0.992 (0.517) | 0.999 (0.516) |
| **Refinement** |  |  |  |
| Resolution range | 52.10-1.50 | 52.17-1.90 | 51.22-1.49 |
| Number of reflections | 28,277 | 11,148 | 29,757 |
| $R_{\text{work}}$ | 17.225 | 20.716 | 18.931 |
| $R_{\text{free}}$ | 19.303 | 24.867 | 21.586 |
| Wilson B-factor ($\text{\AA}^2$) | 16.01 | 19.47 | 18.80 |
| Mean B-factor ($\text{\AA}^2$) | | | |
| Overall | 21.74 | 24.55 | 22.28 |
| Protein | 20.35 | 24.41 | 21.80 |
| Water | 22.20 | 26.64 | 29.70 |
| Ligands | - | 40.76 | 19.31 |
| Other | - | - | 26.22 |
| **Model quality** |  |  |  |
| *RMS deviations* |  |  |  |
| Bond length ($\text{\AA}$) | 0.0103 | 0.0101 | 0.0110 |
| Bond angles ($^\circ$) | 1.70 | 1.81 | 1.75 |

## Notes

### Competing Interest Statement

The authors have declared no competing interest.

